# Correlates of non-random patterns of capsule switching in pneumococcus

**DOI:** 10.1101/811406

**Authors:** Shreyas S. Joshi, M. A. Al-Mamun, Daniel M. Weinberger

## Abstract

**Background:** Pneumococcus is a diverse pathogen, with >90 serotypes, each of which has a distinct polysaccharide capsule. Pneumococci can switch capsules, evading vaccine pressure. Certain serotype pairs are more likely to switch, but the drivers of these patterns are not well understood.

**Methods:** We used the PubMLST and Global Pneumococcal Sequencing (GPS) databases to quantify the number of genetic lineages on which different serotype pairs co-occur. We also quantified the genetic diversity of each serotype. Regression models evaluated the relationship between shared polysaccharide structural components and the frequency of serotype switching and diversity.

**Results:** A number of serotype pairs co-occurred on the same genetic lineage more commonly than expected. Co-occurrence of between-serogroup pairs was more common when both serotypes had glucose as a component of the capsule (and, potentially, glucuronic acid). Diversity also varied markedly by serotype and was lower for serotypes with glucuronic acid in the capsule and higher for those with galactose in the capsule.

**Conclusions:** Certain pairs of serotypes are more likely to occur on the same genetic background, and these patterns were correlated with shared polysaccharide components. This might indicate adaptation of strains to produce capsules with particular characteristics.

## INTRODUCTION

*Streptococcus pneumoniae* (pneumococcus) is commonly found in the upper respiratory tract of healthy children and also causes a large burden of disease in children and adults. Pneumococci are grouped into serotypes based on the structure and antigenic properties of the extracellular polysaccharide capsule [1]. More than 90 serotypes have been identified, with a fraction of these serotypes causing the majority of invasive pneumococcal disease (IPD) cases worldwide.

The capsule itself is the target of pneumococcal conjugate vaccines (PCV). Currently-available conjugate vaccines target the capsules of 10 or 13 of these serotypes [2–4]. These vaccines have driven down rates of severe disease and have also disrupted transmission of vaccine-targeted serotypes among healthy children [5,6]. Because of this disruption to the bacterial ecology, serotype replacement is observed, wherein serotypes not targeted by the vaccine increase in frequency among healthy carriers and, to a lesser degree, as causes of disease [7]. With next-generation vaccines under development that target additional serotypes, it is important to understand the factors that shape the pneumococcal population and the emergence of new strains.

The non-vaccine serotypes that emerge following vaccine introduction result from the tremendous diversity of pneumococcus. Beyond variation in the capsular polysaccharide, only about 70% of the genome is conserved across all pneumococcal strains, and there is a large complement of accessory genes [8,9]. Pneumococci can be classified into genetic lineages using various classification schemes, and genetic lineages can acquire different capsules [10–12]. In a serotype-switching event, a strain producing a particular capsule acquires the genetic cassette required to synthesize a different capsular polysaccharide from another strain [13–16].

Recombination of these loci leads to the production of a different polysaccharide capsule or even the emergence of a novel serotype [17]. These capsule-switching events generate a pool of diversity that can contribute to serotype replacement. In particular, when a genetic lineage that is predominantly expressing a vaccine-targeted capsule (a vaccine serotype, VT) has undergone capsule switching, then a variant of that lineage that expresses a capsule not targeted by the vaccine can increase in frequency [12].

Not all serotype switches are equally likely to occur. A strain is more likely to switch to a structurally-similar serotype in the same serogroup (e.g., between serotypes 19A and 19F), than to a totally distinct serotype in a different serogroups [16,18–21]. Several mechanisms have been proposed to explain these non-random patterns, including genetic similarity of the loci and epistatic interactions between the capsule cassette and the genetic background, but the reasons for these non-random patterns have not been resolved [21].

In this study, we used data from several large global databases of pneumococcal clinical isolates with information on genetic lineage and serotype to evaluate the degree of capsular switching by individual serotypes and to characterize within- and between- serotype switching patterns. We accomplished this by analyzing the frequency of genotype/serotype co-occurrence using several complementary analytic tools. We then considered the potential role of shared biochemical characteristics of the capsules in influencing these patterns.

### Data Sources

PubMLST (https://pubmlst.org/) is a large global database of pneumococcal clinical isolates with information about MLSTs. Allele and MLST assignments for *Streptococcus pneumoniae* isolates in the PubMLST database were used in this analysis [18][22]. Isolates for which the serotypes were non-typeable or with errors in their names were excluded from the analysis. The final dataset consisted of 35,898 isolates, representing 96 serotypes and 11,718 MLST lineages from the data available on March 23, 2019. Based on the allelic similarity between isolates, 581 clonal clusters (CC) were generated using the eBURST [23] tool from the PubMLST database. As a comparison, we analyzed the serotype and sequence cluster (GPSC) data for 13,454 isolates from the Global Pneumococcal Sequencing Project (GPS) presented by Gladstone et al [24]. Throughout the manuscript, ‘genetic lineage’ refers to either the MLST, CC, or GPSC group, as indicated. We used polysaccharide composition data from the previously published literature [25], and the carriage data were as described in Tothpal et al [26].

### Detecting non-random serotype-switching patterns

We assumed that if two serotypes were detected on the same genetic lineage, this resulted from a capsule switching event. We used two complementary methods to detect such capsule switching events: Monte Carlo (MC) [18,27,28] and market basket (MB) [29,30] analysis (also called Association Rule Mining). We used MC simulation to assess if the number of genetic lineages associated with a serotype pair were more than what was expected by chance. We first counted the number of genetic lineages on which a serotype pair was detected, i.e., the observed number of shared genetic lineages. These pairs were then reshuffled to produce 1000 random datasets, and for each pair, the number of shared genetic lineages were calculated. 97.5 percentile was chosen as a significance threshold, and serotype pairs with shared genetic lineages higher than this threshold were considered significant candidates for potential serotype switching.

As a complementary analysis, we used MB analysis, which is used in commerce settings to identify associations between items likely to be purchased together. In our analysis, an individual genetic lineage is like a customer shopping for different capsule structures or serotypes. For a serotype pair AB among N total pairs, ‘support’ represents the proportion of all pairs that contain AB *frq(A,B)/N*. Confidence is calculated as *frq(A,B)/frq(A)* and represents the proportion of times that B is detected when A is detected. And the lift is calculated as *Support(AB)/(Support(A)*\**Support(B))* with values >1 indicating how much more the pair occurs than would be expected if they were independent. Significance values for Chi-square test were calculated to assess dependence between serotypes in a pair, and Fisher’s exact test to measure the significance of association rules.

Aggregating the data by MLST and CC has tradeoffs for detecting serotype switch events. The patterns observed can vary considerably for MLST and CC. To assign overall importance values to serotype associations from the MB and MC analysis results, we built an index by assigning values to the output of the different statistical analyses of our data and then summing them. Serotype pairs with frequency/count of 3 or more were assigned a value of 1. For Monte Carlo output, serotype-pairs were assigned a value of 1 if observed values were greater than the threshold of 97.5 percentile. For Market Basket output, a value of 0.25 was assigned for pairs with confidence measure values in the top quartile, 0.25 if chi-square test is significant, 0.25 for lift >1, and 0.25 if fisher’s exact test is significant. Essentially we have assigned values of 1 to the count, 1 to Monte Carlo, and 1 to Market Basket (maximum value of 3). The values for both MLST and CC are then combined, thus giving a range of index scores from 0 to 6, with larger scores indicating stronger evidence of a link. Based on the index output, serotype associations were visualized via a network plot. Consensus network plots were built for pairs with index score thresholds from 1 to 5, with more or fewer connections shown in the plot (supplementary figure). The network for serotype pairs with index scores greater than 4 provided the best balance of complexity ability to visually interpret the plot.

### Serotype diversity

Serotype diversity was calculated based on how many isolates of that serotype were identified in different clonal clusters. We made a Serotype*Genetic Lineage matrix containing the number of isolates for each unique combination of serotype and genetic lineage. Simpson’s Diversity Index (SDI) is a tool used in population genetics to measure diversity within populations.[31,32] We used it to measure the diversity of the genetic lineage within individual serotypes. In this case, the SDI value represents measure of probability that in a sample where all isolates belong to the same serotype, two randomly selected isolates have different genetic lineages. SDI were calculated for each serotype using the formula Σn(n−1)/N(N−1),where N is the total of isolates belonging to one serotype and n is the number of isolates for a single genetic lineage within a serotype.

### Correlation of capsule switch patterns and diversity with polysaccharide characteristics

We hypothesized that the inclusion of specific sugars in the capsule could influence the frequency of capsule exachange. We therefore evaluated correlations between specific sugars and the co-occurrence of different serotypes on the same genetic lineage. To evaluate the correlation between capsule exchange patterns and capsule structure, we performed a series of regression analyses. The goal was to test whether serotypes that had sugar X in their capsule were more likely to switch with another serotype that also had sugar X in its capsule. For every potential pair of serotypes, we counted the number of genetic lineages that they were both detected on. We then fit a series of quasi-Poisson regressions in which the outcome variable was the number of genetic lineages on which both serotypes were detected. The main covariate of interest was whether sugar X was present in both members of the serotype pair or just 1 member of the pair. A positive coefficient would indicate that the pairs in which both members have sugar X in the capsule are more common than serotype pairs in which only one member of the pair has sugar X. Serotypes found on more genetic lineages would be more likely to co-occur with each other by chance. Therefore, we adjusted for this in the regression by using the product of the proportion of genetic lineages on which serotype A was found and the proportion of genetic lineages on which serotype B was found. This product was normalized using a Box-Cox transformation (lambda=0.01). Each sugar was tested 1 at a time. The analyses were restricted to serotype pairs that were not in the same serogroup and in which at least 1 member of the pair had sugar X in the capsule. The analyses were repeated separately when grouping strains by MLST or by CC. We also performed a similar set of analyses but where the outcome was binary (does the serotype pair co-occur on any MLST/CC or not?). This was analyzed using a logistic regression model. The results from all 4 sets of analyses (CC/MLST, quasi-Poisson/logistic) are presented together for comparison.

As a complementary analysis, we evaluated the correlation between the presence of specific sugars in the capsule and the serotype-specific SDI. To evauate the correlation between polysaccharide components and SDI, we performed linear regression, where the outcome variable was the serotype-specific SDI (logit-transformed). We expected that SDI would be associated with the prevalence of the serotype (because more commons serotypes have more opportunity for recombination), so we controlled for carriage prevalence in the pre-vaccine period, as measured in the UK by Sleeman et al. [26,33]. We then tested whether the presence or absence of individual sugars in different serotypes was associated with SDI. 13 out of 25 sugars were present in the capsules of 3 or more serotypes and were included in the analysis. Each sugar was tested individually in a series of regression models; each model included covariates for carriage prevalence and the presence/absence of 1 sugar at a time. Akaike information criterion (AIC) values and regression coefficients with 95% confidence intervals have been reported for individual sugars.

The analyses were performed in Rstudio IDE[34] for programming in R [35]. Custom scripts were written for SDI, MC, and scraping the eBURST output. The ‘arules’ package was used for performing MB [36]. The networks were generated using packages ‘igraph’[37] and ‘ggplot2’ [38]. The output of igraph was saved in graphml file which was then visualized in Cytoscape [39]. Phylogenies based on polysaccharide composition and figure of tree was made using ‘ape’ and ‘phytools’ packages respectively. The code and data used for the analysis is available at https://github.com/weinbergerlab/sero_switch_paper.

## RESULTS

### Non-random patterns of within and between serogroup switching

We quantified the degree of serotype switching, both within and between serogroups, for every possible pair of serotypes. In agreement with previous studies [18], we found a higher-than-expected likelihood of serotype switching within serogroups (**Figure 1**). A number of outside-serogroup switches were also detected more frequently than expected by chance.

**Figure 1.**
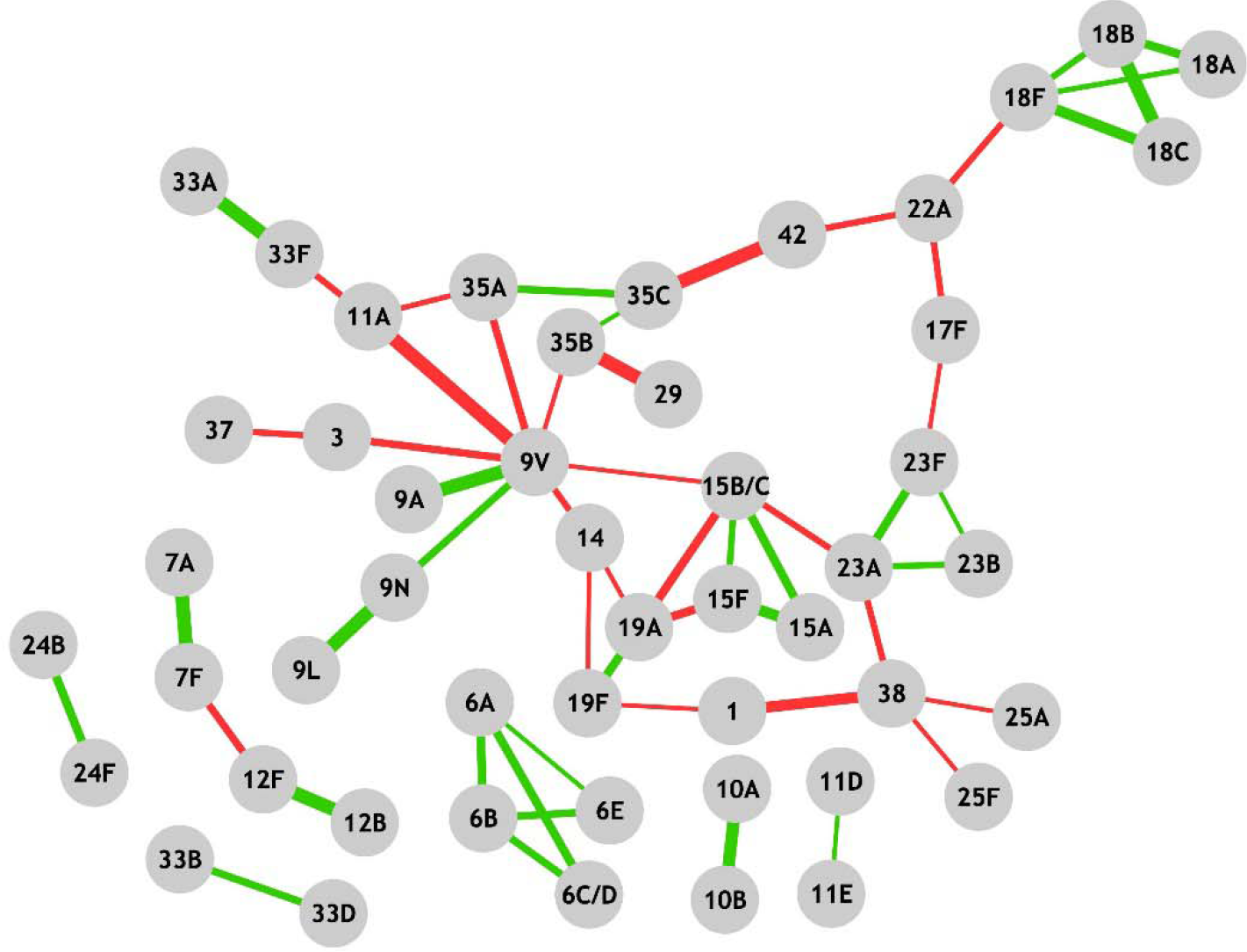
Consensus network of serotype associations. Pairs with index scores of 4 or greater are shown. The width of the edges indicates index scores. Green edges indicate within serotype switches, red edges indicate between serotype switches.

We hypothesized that serotype pairs with capsular polysaccharides that are more similar biochemically might be more likely to switch. Serotype pairs in which both members had glucose in their capsules were more likely to be detected on the same genetic background (comparing between-serogroup pairs only) (**Table 1**). This result was consistent whether aggregating the isolates by CC, MLST, or GPSC and was confirmed by examining the top co-occurring serotype pairs from the market basket/Monte Carlo analyses (**Table 2)**. Glc was found in the structure of both serotypes in all but one of the pairings for which structural information was available (overall, Glc is present in the capsules of 67 of 84 serotypes with known structure). There was also some evidence that serotypes that had GlcA in the capsule were more likely to co-occur on the same genetic lineage as another serotype with GlcA (**Table 1**); however there was a high degree of uncertainty in the estimates when the isolates were grouped by MLST or GPSC rather than CC. For other sugars (e.g., FucNAc, any NAc sugar), there were some associations depending on how the strains were grouped or analyzed, but the patterns were not consistent (**Table 1)**. There were not notable similarities between clusters identified based on overall similarity of polysaccharide structures and the between-serogroup pairs identified in the network plot (Supplementary figure: phylo_tree).

**Table 1.** Association between the presence of specific sugars in the capsule and co-occurrence of serotype pairs on the same genetic background (MLST, CC, or GPSC).

|  | **Poisson MLST** | **Logistic MLST** | **Poisson.CC** | **Logistic.CC** | **Poisson GPSC** | **Logistic GPSC** |
| --- | --- | --- | --- | --- | --- | --- |
| Glc | 0.16 (0.02,0.29) | 0.3 (0.09,0.51) | 0.13 (0.08,0.17) | 0.16 (0.01,0.3) | 0.5 (0.33,0.68) | 0.43 (0.22,0.64) |
| GlcA | 0.16 (-0.18,0.46) | 0.14 (-0.45,0.71) | 0.18 (0.04,0.32) | 0.09 (-0.43,0.61) | 0.19 (-0.32,0.64) | 0.46 (-0.16,1.06) |
| Any NAc | 0.43 (0.32,0.53) | 0.39 (0.15,0.63) | 0.01 (-0.04,0.06) | -0.12 (-0.28,0.04) | 0.42 (0.28,0.56) | 0.45 (0.23,0.68) |
| Rha | -0.24 (-0.36,-0.12) | -0.41 (-0.72,-0.1) | -0.01 (-0.07,0.05) | 0.28 (0.06,0.51) | -0.27 (-0.44,-0.1) | -0.56 (-0.85,-0.27) |
| Gal | -0.21 (-0.32,-0.1) | 0.06 (-0.14,0.26) | 0.06 (0.02,0.11) | 0.41 (0.27,0.56) | -0.17 (-0.3,-0.04) | 0.08 (-0.12,0.27) |
| GlcNAc | -0.05 (-0.35,0.23) | -0.29 (-0.85,0.24) | 0.12 (0.01,0.22) | -0.03 (-0.35,0.28) | 0.39 (0.1,0.66) | 0.54 (0.15,0.93) |
| Gro | -0.18 (-0.48,0.1) | -0.39 (-1,0.2) | 0.11 (0,0.21) | 0.45 (0,0.9) | 0.36 (0.12,0.59) | 0.26 (-0.22,0.72) |
| FucNAc | 1.33 (0.72,1.85) | 2.67 (1.18,4.17) | -0.02 (-0.4,0.33) | -0.31 (-1.41,0.74) | 1.57 (0.3,2.54) | 1.89 (0.54,3.06) |
| Rib-ol | 0.13 (-0.13,0.38) | 0.31 (-0.09,0.69) | -0.03 (-0.12,0.06) | 0.25 (-0.01,0.51) | 0.6 (0.39,0.81) | 1.59 (1.22,1.95) |
| ManNAc | 0.37 (0.15,0.57) | 1.18 (0.32,2.06) | -0.03 (-0.17,0.1) | 15.93 (-4.33,178.85) | 0.19 (-0.23,0.56) | 0.28 (-0.48,1.03) |

**Table 2.** Pairs of serotypes found together on the same genetic background more commonly than expected, and the polysaccharide components shared between them.

| Serotype1 | Serotype2 | Index score | Shared polysaccharide components |
| --- | --- | --- | --- |
| 11A | 9V | 6 | Ac, Gal, Glc |
| 29 | 35B | 6 | Gal, GalNAc, Rib-ol |
| 35C | 42 | 6 | Gal, Glc, Man-ol |
| 1 | 38 | 5.5 | 38* |
| 15B/C | 19A | 5 | Glc |
| 15F | 19A | 5 | 15F* |
| 12F | 7F | 4.75 | Gal, GalNAc, Glc |
| 3 | 9V | 4.75 | Glc, GlcA |
| 35A | 9V | 4.75 | Ac, Gal, Glc |
| 11A | 33F | 4.5 | Ac, Gal, Glc |
| 14 | 9V | 4.5 | Gal, Glc |
| 15B/C | 23A | 4.5 | 23A* |
| 17F | 22A | 4.5 | 22A* |
| 18F | 22A | 4.5 | 22A* |
| 22A | 42 | 4.5 | 22A* |
| 23A | 38 | 4.5 | 38* |
| 3 | 37 | 4.5 | Glc |
| 11A | 35A | 4.25 | Ac, Gal, Glc |
| 1 | 19F | 4 | NA |
| 14 | 19A | 4 | Glc |
| 14 | 19F | 4 | Glc |
| 15B/C | 9V | 4 | Ac, Gal, Glc |
| 17F | 23F | 4 | Gal, Glc |
| 25A | 38 | 4 | 25A*, 38* |
| 25F | 38 | 4 | 38* |
| 35B | 9V | 4 | Ac, Gal, Glc |

Between serogroup pairs with index scores of 4 or more. Shared polysaccharide components contain the sugars shared between the serotypes in a pair. Polysaccharide composition data was not available for serotypes 38, 23A, 25A, 22A and 15F. Abbreviations of sugars are Ac, acetate; Gal, galactose; GalNAc, N-acetylgalactosamine; Glc, Glucose; GlcA, glucuronic acid; Man-ol, mannitol; Rib-ol, ribitol.

### Variation in diversity by serotype

Another measure of the amount of capsule switching occurring for each serotype is based on the diversity of clonal clusters associated with a serotype—serotypes that switch more readily will be associated with more genetic lineages. There was considerable variability in diversity between serotypes **(Figure 2)**. We considered whether the presence of certain sugars in the capsule was associated with higher or lower diversity. The hypothesis is that the inclusion of certain sugars in the capsule could influence the probability that the capsule could efficiently switch to a different genetic lineage, due to epistatic interactions. In regression analysis, after controlling for pre-vaccine carriage frequency, diversity was lower for serotypes with GlcA in their capsules and higher for serotypes with Gal in the capsule (**Table 3**). The association between GlcA and diversity was apparent when analyzing the data by CC or by GPSC, but not when aggregating by MLST. The association with Gal was similar regardless of grouping, but when grouping by GPSC there was more uncertainty in the estimate.

**Table 3.** Association between the diversity of a serotype and the presence of specific sugars in the capsule, with diversity based on MLST, CC, or GPSC

|  | **Coefficient (CC)** | **Coefficient (MLST)** | **Coefficient (GPSC)** |
| --- | --- | --- | --- |
| Gal | 0.91, (0.31,1.52) | 0.65, (0.15,1.15) | 0.63, (-0.3,1.56) |
| GlcA | -1.05, (-1.75,-0.35) | -0.12, (-0.74,0.51) | -1.75, (-2.7,-0.79) |
| Gro | 0.65, (-0.07,1.36) | 0.18, (-0.43,0.79) | 0.74, (-0.29,1.77) |
| GlcNAc | 0.43, (-0.27,1.12) | -0.2, (-0.77,0.37) | 0.47, (-0.47,1.41) |
| Ac | -0.26, (-0.86,0.33) | -0.37, (-0.84,0.1) | 0.05, (-0.8,0.9) |
| FucNAc | -0.21, (-1.23,0.81) | -0.34, (-1.16,0.47) | -1.39, (-2.89,0.1) |
| ManNAc | 0.16, (-0.68,1) | 0.46, (-0.26,1.18) | 0.52, (-0.7,1.73) |
| GalNAc | -0.07, (-0.76,0.62) | -0.47, (-1.02,0.07) | -0.03, (-1.06,1) |
| Glc | 0.1, (-0.61,0.8) | -0.18, (-0.75,0.39) | 0.23, (-0.79,1.26) |
| Rha | -0.01, (-0.61,0.6) | 0.25, (-0.24,0.74) | 0.14, (-0.72,0.99) |

**Figure 2:**
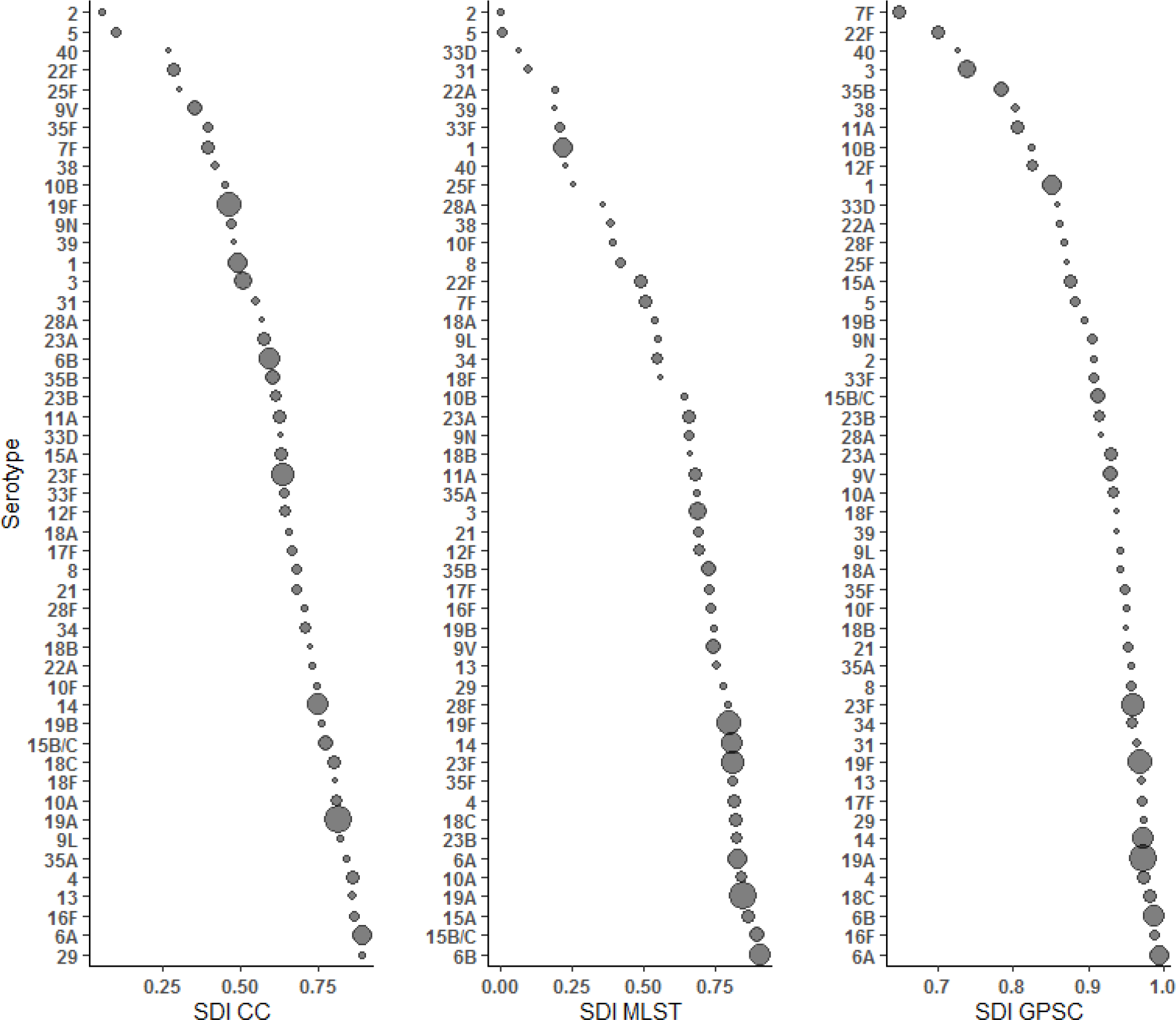
Simpson’s Diversity Index for each serotype, based on the diversity of associated clonal clusters (CC), multi-locus sequence types (MLST), or Global Pneumococcal Sequence Clusters (GPSC). The size of the bubble is propotional to the number of isolates of the serotype in he respective databases.

## DISCUSSION

In this study, we demonstrate substantial variability in genetic diversity between serotypes. Variation in diversity was correlated with the presence of specific components in the polysaccharide capsule (GlcA and Gal). And non-random capsule switch events were detected, both within and between serogroups. Switching between serogroups was more common when both members of the pair had Glc (or possibly GlcA) in the capsule structure. Together, these results support the notion that there is an interaction between genetic background and the capsule. Some capsules had more or less ability to switch to a different genetic background, and some genetic backgrounds could be well-adapted to produce capsules with particular characteristics. We hypothesized that the observed serotype-genotype patterns could be driven by the sugar composition of the capsule itself due to epistatic interactions between capsular genes and components of central metabolism [40]. In this scenario, a successful capsule switch event would be the one that results in an antigenic shift in the capsule structure but that would have similar metabolic properties. Metabolic tradeoffs could explain why serotypes with GlcA in the capsule were less diverse on average—if other non-capsular factors are needed to efficiently produce a capsule containing GlcA, there would be a lower probability of switching. GlcA is found in the capsules of only 13 of the 84 serotypes that have known polysaccharide structures and is associated with lower diversity values. Some strains could more efficiently respond to the metabolic flux that results from shunting UDP-Glc into UDP-GlcA for capsule production [41]. Or alternatively, strains could differ in their ability to uptake and use hyaluronic acid (HA). HA is a disaccharide glycoconjugate with GlcA and N-acetylglucosamine (GlcNac) as its components [42] and has been reported to be used by pneumococcus as a sole carbon source for growth [43]. Hyaluronate lyase released by pneumococcus can break down HA into disaccharide units.[44] If pathways exist in some strains to incorporate this exogenous GlcA into the capsule (via conversion to UDP-GlcA), it could provide a fitness advantage. Experimental studies would be needed to determine the mechanism underlying these patterns.

Aside from metabolic explanations, serotype switching patterns could also be influenced by the structure of the capsular biosynthesis locus, which might influence the probability of having a successful homologous recombination event [21]; or switching patterns could be influenced by other characteristics of the serotypes that influence the uptake or exchange of DNA. And finally, the mechanism of initiation of polysaccharide chain formation could influence capsule switching. Most serotypes initiate their polysaccharide chain with glucose [45]. Characteristics of the cell surface on certain strains could be better adapted to link to glucose than to other possible initiating sugars.

Our analyses have certain limitations. We cannot determine the direction of the serotype switch from the MLST data, which would be possible with more detailed WGS data[46]. *Streptococcus pneumoniae* is one of the first organisms to have a critical mass of whole genome sequence (WGS) data that can potentially assist with studying genetic mechanisms of serotype switching. There are now tens of thousands of pneumococcal genomes that have been sequenced [47][24] [14–16], providing an opportunity to further interrogate some of the putative serotype switching patterns we have identified here. The reliance on the MLST database has other limitations as well. For instance, we were not able to verify the validity of the serotype assignments, and for certain pairs of serotypes (29/35B), the association could be due to misclassification of the serotype by the original investigators. These two serotypes are structurally similar and can sometimes be difficult to distinguish with routine methods. However, the data from the GPS project, which are based entirely on DNA sequence rather than laboratory-based serotyping, affirmed the main results of our analyses. Finally, the analyses presented here are based on correlations using genetic data. Experimental work is needed to further understand the barriers to capsule switching and the potential importance of specific components of the capsule in influencing the patterns of capsule switching.

In conclusion, we use information on the co-occurrence of serotypes on the same genetic background to evaluate capsule switching patterns in pneumococcus and to explore possible barriers to capsule switching. The associations of serotype diversity and specific capsular components suggests that there could be important interactions between the capsule and the genetic background on which the capsule is expressed. Such interactions could help to shape how pneumococci respond to vaccine-related selective pressure. This information could help to predict which serotypes are most likely to emerge in the future.

---

**Supplementary Figure:**
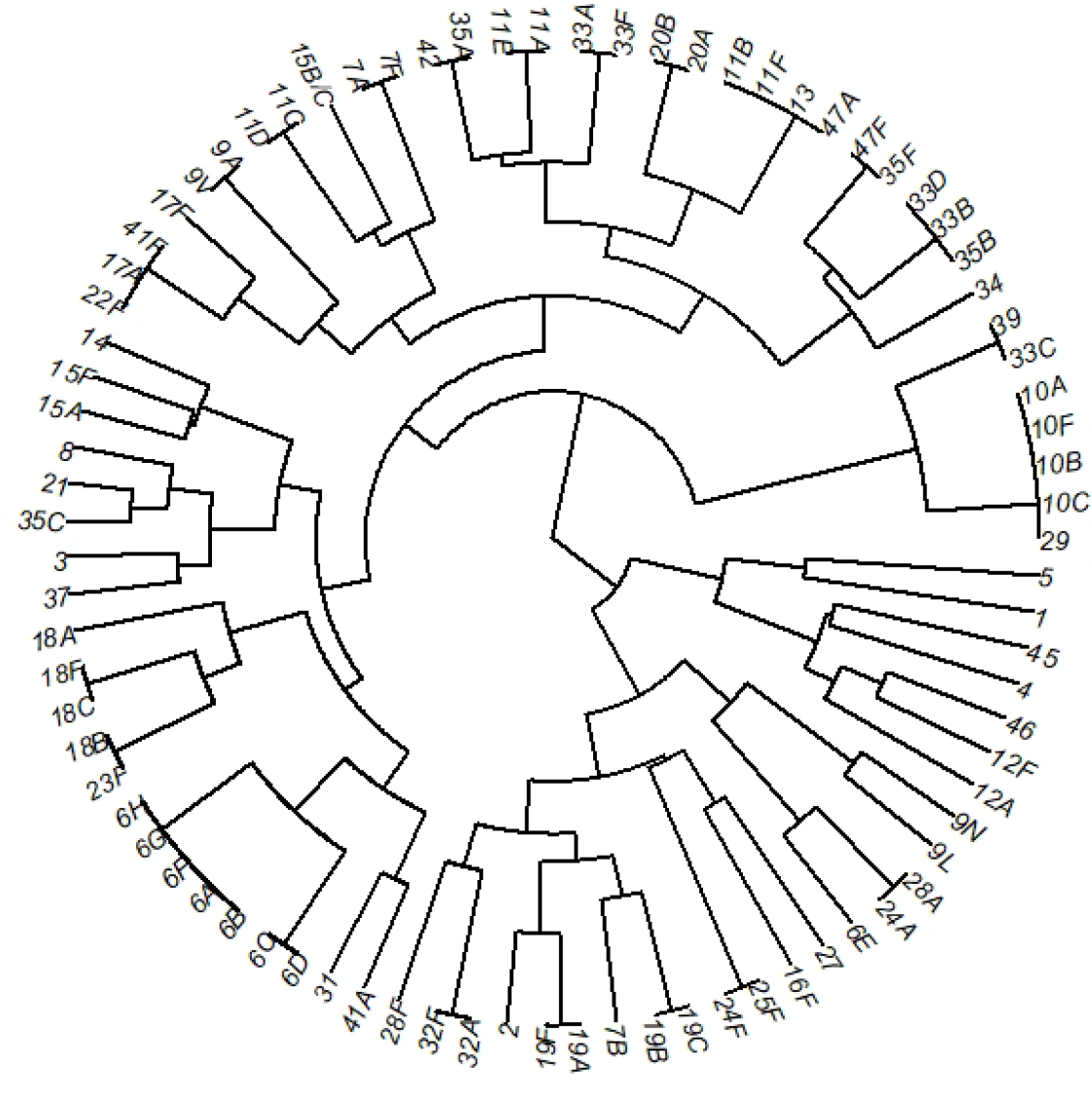
Phylogenetic tree output for clustering based on the polysaccharide composition in serotype capsules.

